# Closed-Loop Vibrotactile Neuromodulation for Reducing Tremor-Related Propranolol Use

**DOI:** 10.64898/2026.08.07.743626

**Authors:** Varun Agarwal, Arin Soneji

## Abstract

Pathological tremor is a neurological condition that impairs fine motor tasks, affecting 1% of the general population and 4% of the elderly. Tremors arise when muscles’ micro-oscillations synchronize and phase lock, typically within a 4–12 Hz frequency range. Administering beta-blockers can reduce tremor severity, but doses are hard to personalize, with heavy doses of propranolol correlating with low blood pressure, dizziness, and nausea. In this project, we aimed to model tremor and create a closed-loop control framework to suppress tremor amplitude while minimizing pharmacological dependence. Because side effects constrain the use of pharmacological suppression alone, we investigated noninvasive neuromodulation. We used vibrotactile stimulation (VTS) to disrupt pathological tremor synchronization and reduce oscillatory amplitude. We hypothesized that tremor suppression involving VTS followed a nonmonotonic relationship, tested by determining whether maximum relief requires an adaptable framework. The procedure consisted of constructing a propranolol-reduction simulation by implementing a Hill curve, where we calculated and utilized tremor reduction, heart rate (HR) drop, and blood pressure (BP) drop. We then built a device to capture tremor-related data and create vibration using two linear resonant actuator (LRA) coin motors. We connected it to a microcontroller, where we determined optimal vibration frequencies through a feedback loop. Across 50 trials, VTS alone reduced tremor amplitude by an average of 37.3%, reducing the propranolol dose needed to reach 50% total tremor reduction by 71.9%, lowering the modeled blood pressure drop from 38.1 to 18.9 mmHg. This device demonstrates proof-of-concept for a nonmonotonic tremor-vibration relationship to reduce dependency on propranolol in the treatment of pathological tremor. These propranolol dose-reduction estimates are derived from computational simulation and have not been clinically validated; they are not intended as a recommendation to alter prescribed medication.

## INTRODUCTION

Pathological tremor is the world’s most common movement disorder, affecting 1% of the general population and 4% of the elderly. Tremors arise when muscles’ micro-oscillations synchronize and phase lock, typically within a 4–12 Hz frequency range (1,2). These synchronized oscillations normally come from unusual cerebro-thalamo-cortical circuit activity (1). Dopamine is a primary neurotransmitter in aiding motor coordination. A deficiency in this neurotransmitter can cause unintended and chronic movement. Stress, particularly through the release of epinephrine and adrenaline, can also worsen pathological tremor and can independently produce stress-induced tremor due to the increasing destabilization of the central nervous system (3,4). Beta-blockers are competitive antagonists of adrenaline and epinephrine and bind to beta-1 and beta-2 cell receptors, acting as a non-selective inhibitor to these amplifying neurotransmitters (5).

Clinicians favor propranolol for tremor treatment because it binds to and blocks beta-2 receptors, which not only decreases spindle gain but also suppresses oscillatory motions (6,7,8,9). However, propranolol dosing is difficult to personalize and limit, and heavy doses correlate with low blood pressure, dizziness, and nausea (5,10,11). The relationship between propranolol dose and tremor reduction follows a negatively accelerated path, so patients require larger and larger doses to achieve smaller increments of reductions in tremor amplitude (8,9).

Because these side effects limit pharmacological suppression alone and stress-suppression feasibility, researchers have investigated noninvasive neuromodulation as an alternative (12,13). Vibrotactile stimulation (VTS) is one of the approaches, as studies have proven VTS effective for both pathological and stress-induced tremor (13,14,15,16,17,18). Its functionality involves injecting noise into phase-locked, synchronized muscle oscillations to destabilize these waves and reduce tremor (16,17). We hypothesized that tremor behaves like a nonlinear oscillatory system, as VTS only disrupts tremor effectively within specific parameter ranges (and potentially enhances tremor in others), producing a non-monotonic relationship between tremor amplitude and optimal stimulation intensity (17,18,19).

We tested this hypothesis by continuously adapting vibration therapy based on real-time tremor sensing to quantify maximum relief and reduction in dosage ramifications in comparison to constant VTS. In this project, we modeled tremor and developed a closed-loop control framework to measure tremor amplitude suppression and pharmacological dependence minimization. We optimized vibrotactile stimulation in a regression framework to determine optimal neuromodulation frequency and amplitude (Figure 1). Across 50 trials, constant Hz VTS reduced tremor amplitude by an average of 37.3% alone. However, when implementing a closed-loop framework, user-adaptable VTS reduced tremor amplitude by an average of 48.78%. When we paired adaptable VTS with propranolol, the dosage needed to reach half the total tremor reduction fell from 320 mg to 90 mg, a 71.9% reduction. This dropped the blood pressure ramifications from 38.1 to 18.9 mmHg. These findings suggest that adaptable vibrotactile neuromodulation offers a viable strategy for reducing reliance on beta-blockers and mitigating associated clinical side effects in comparison with constant VTS.

**Figure 1.**
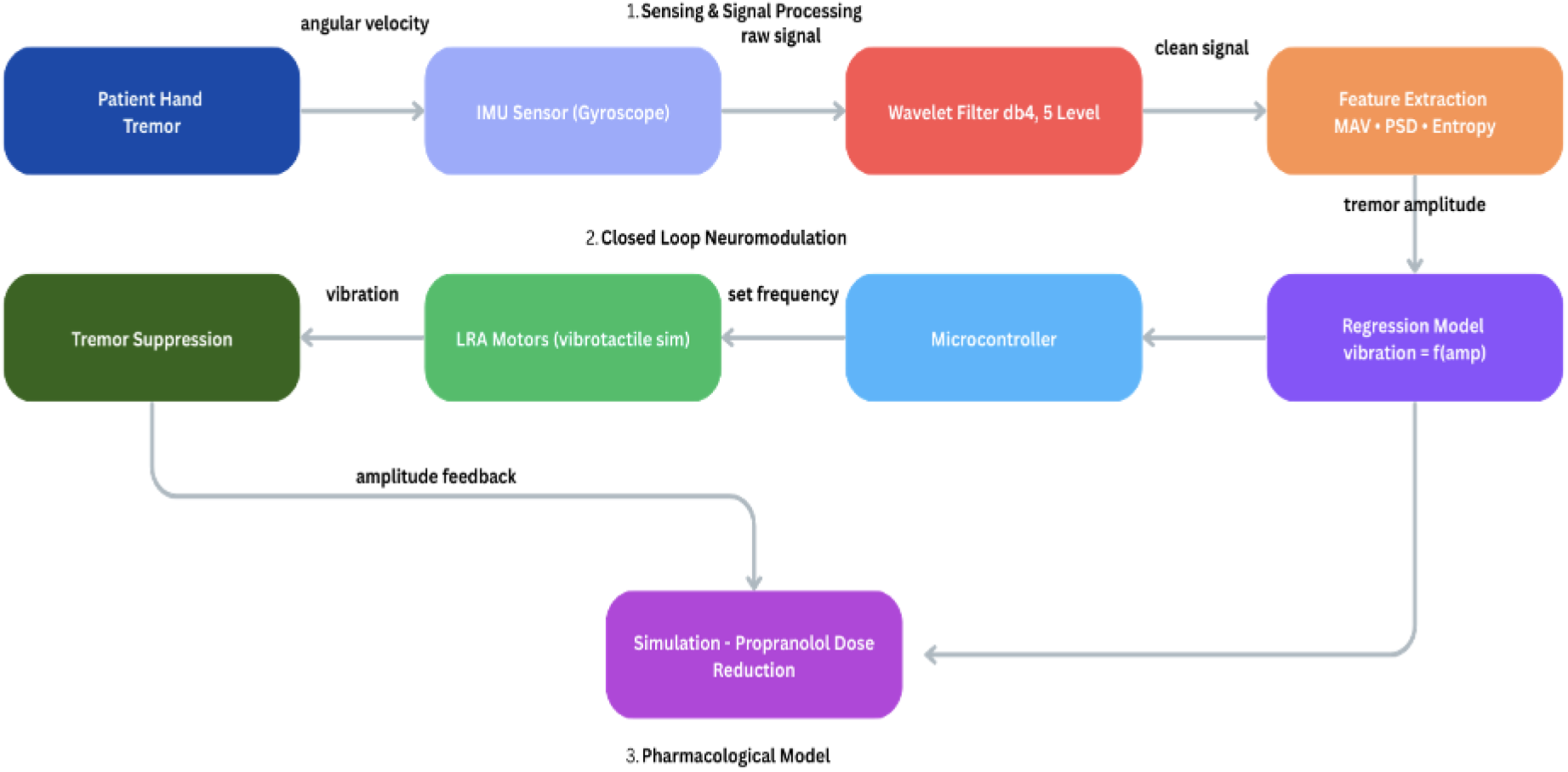
System architecture of the closed-loop vibrotactile neuromodulation device. Block diagram showing the three integrated subsystems: (1) sensing and signal processing, where an IMU captures raw angular velocity from hand tremor and a five-level db4 wavelet filter extracts a clean tremor signal for feature extraction (mean absolute value, power spectral density, approximate entropy); (2) closed-loop neuromodulation, where a regression model maps tremor amplitude to an optimal vibration frequency delivered by two LRA motors, with real-time amplitude feedback; and (3) the pharmacological model, which uses closed-loop suppression output to inform the simulated propranolol dose-reduction calculation.

The goal of the following prototype is to construct a synergistic closed-loop vibrotactile neuromodulation device to minimize dosimetric propranolol modeling for pathological and stress-induced tremor mitigation, confirming this nonmonotonic relationship. The construction and evaluation of this device depend on three key components of design: a probabilistic stress-response model, simulating propranolol dosage alongside tremor reduction, and a closed-loop design for optimizing tremor reduction percentage for vibrotactile stimulation. The probabilistic stress-response model takes in physiological data such as heart rate and temperature, and creates a score from 0 to 1 on how stressed an individual is. This score will be utilized in further research quantifying segmented reduction of propranolol. The propranolol dosing simulation measures propranolol dosing and its corresponding tremor amplitude reduction percentage as a baseline measurement (tremor reduction percentage and blood pressure drop) to determine the effectiveness of the optimized vibrotactile stimulation on minimizing dosage & side effects. The device for vibrotactile stimulation will be the component that aims to reduce the side effects of propranolol dosing while maximizing the percentage of tremor reduction. This device will modify the frequency and amplitude of the vibration therapy in real-time, depending on the changing frequency and amplitude of the physiological tremor. Tremor reduction percentage will be mapped accordingly, and results will be compared to the baseline mentioned in the propranolol dosing simulation and stress-response model. Other alternatives previously explored include median and radial nerve stimulation and transcutaneous vagus nerve stimulation.

In order for our prototype to successfully meet our design criteria and accomplish the design goal, each key component will be comprehensively evaluated with different criteria. The probabilistic stress-response model will be evaluated on the model’s AUC (Area Under the Curve). AUC represents the model’s ability to correctly identify stress amounts and states. The propranolol dosage simulation will be evaluated through simulated tremor amplitude reduction percentage and blood pressure drop. The closed-loop vibrotactile stimulation device will be measured through simulated tremor amplitude reduction and translated blood pressure effects (using less propranolol). It will be compared to the simulation baseline to determine efficacy.

## RESULTS

### Stress Classification Model

The stress classification model achieved an AUC of 0.919 (Figure 2), showing strong accuracy in distinguishing stress from non-stress states (3,20). We fed these classifications directly into the closed-loop tremor reduction simulation. This high accuracy allows for reliability in measuring the stress component of suppression. Potential limitations with this model include reporting 807 false negatives and 450 false positives against 6,623 true negatives and 1,186 true positives, yielding a false positive rate of 0.064 and a false negative rate of 0.405. However, this aligns with the design goal of tolerating false negatives over false positives to avoid incorrectly triggering dosage. The most balanced operating point placed recall at 0.629 and precision at 0.712.

**Figure 2.**
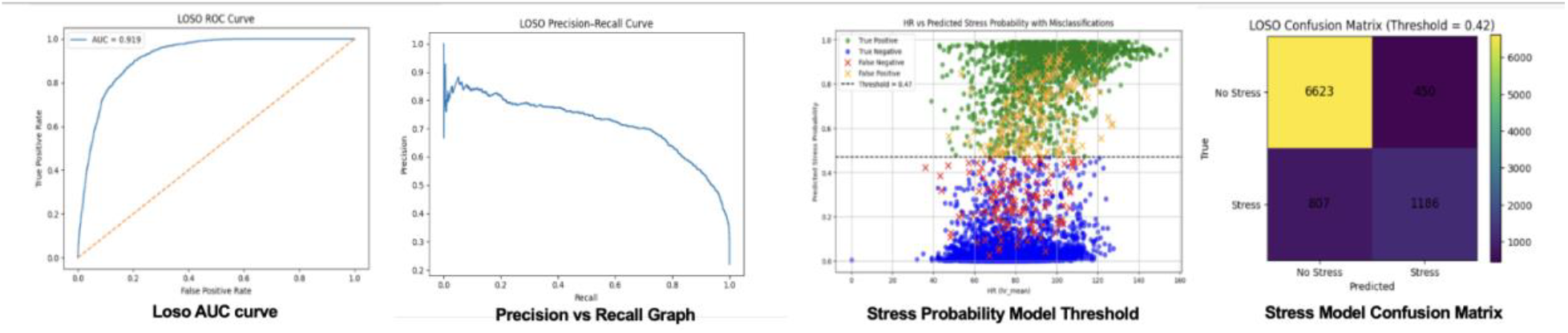
Performance evaluation of the probabilistic stress-response model under leave-one-subject-out (LOSO) cross-validation. (A) ROC curve (AUC = 0.919). (B) Precision-recall curve. (C) Predicted stress probability plotted against mean heart rate, colored by classification outcome relative to the operating threshold. (D) Confusion matrix at the selected classification threshold.

To model tremor reduction from stress, we developed a simulation where stress probability influenced a pharmaceutical response model, using a Hill curve to predict effects on heart rate, tremor, and blood pressure (21). We analyzed many dosing strategies, tracking each of them across benefit, risk, efficiency, severity, and blood pressure impact, then combined them into a single total score. The stepwise strategy performed the best based on the blood-pressure-penalized score, producing roughly a 30% decrease in tremor alongside a 15% drop in blood pressure (Figure 3) (10,11).

**Figure 3.**
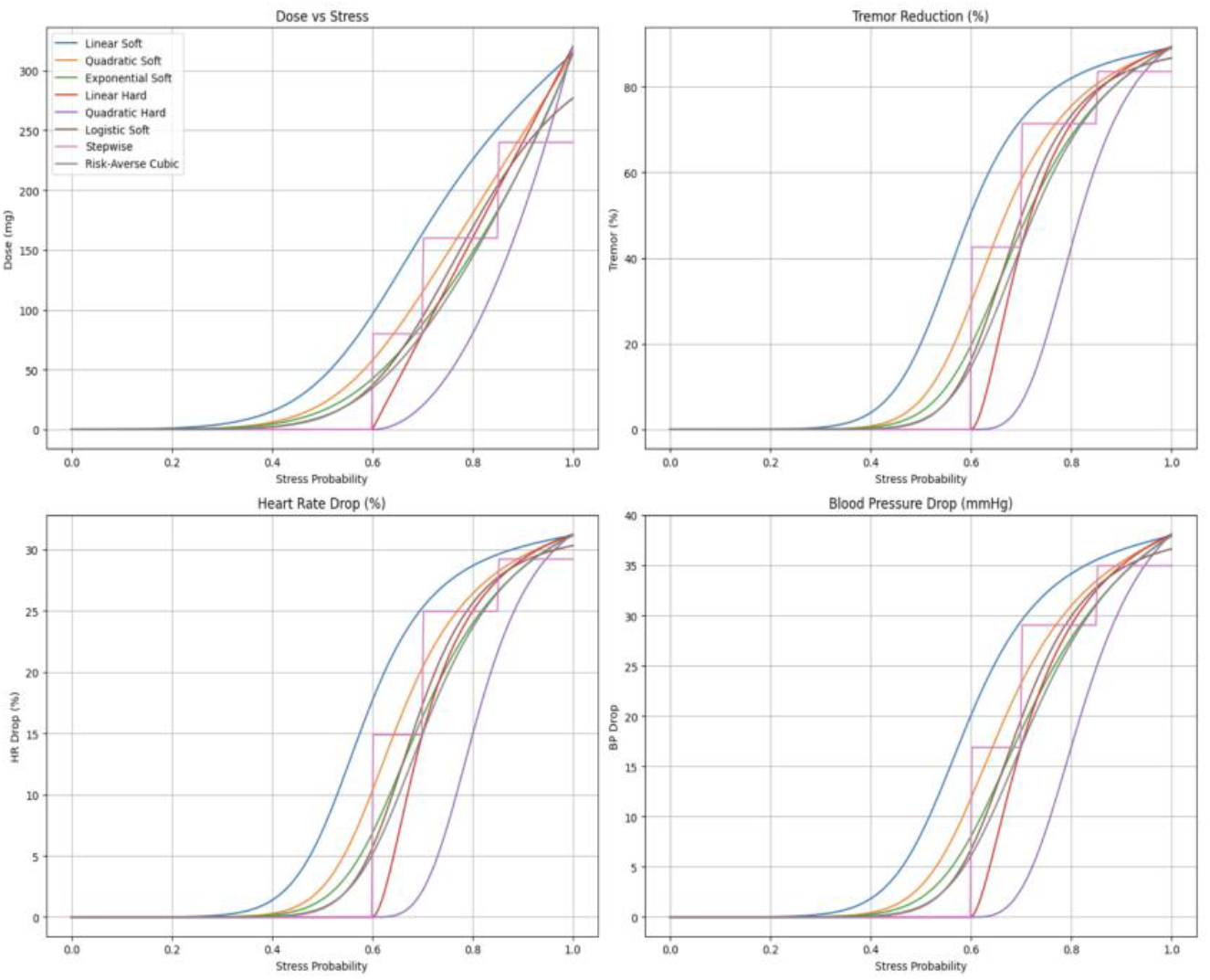
Comparison of candidate propranolol dosing strategies as a function of modeled stress probability. Eight strategies (Linear Soft, Quadratic Soft, Exponential Soft, Linear Hard, Quadratic Hard, Logistic Soft, Stepwise, and Risk-Averse Cubic) are compared across simulated propranolol dose (top left), tremor reduction (top right), heart rate drop (bottom left), and blood pressure drop (bottom right), each plotted against stress probability. The stepwise strategy was selected based on the best blood-pressure-penalized benefit-risk score.

We also trained a Hill equation model to predict tremor reduction based on vibration intensity and tremor amplitude. It predicted an average tremor reduction of 35.7% across all tested conditions, with a peak reduction of 69.0% at a tremor amplitude of 1.47 using an optimal vibration intensity of 114. Observed tremor amplitudes ranged from 1.47 to 3.44, averaging 2.42. The model fit the data well, with an R^2^ of 0.791, following the regression: y = 120x^0.485/(2.407^0.485 + x^0.485) (Figure 4).

**Figure 4.**
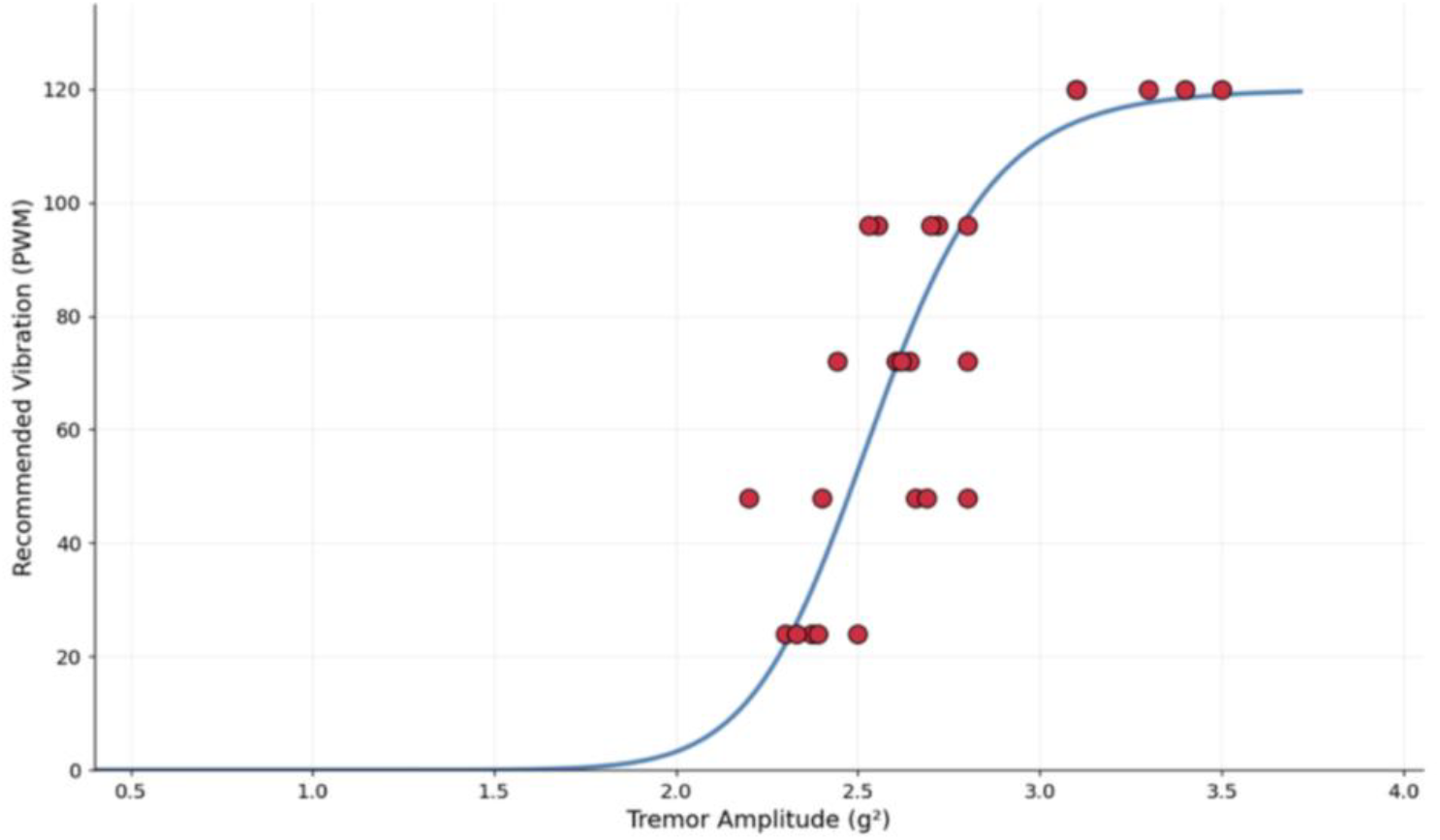
Optimal regression curve for recommended vibration intensity based on tremor amplitude. Dose-response curve showing the recommended vibration intensity across different tremor amplitudes, with dots representing individual data points (n=2). Curve generated using a Hill equation fit. R^2^ = 0.791.

Constant vibrotactile stimulation alone achieved 37.3% tremor suppression, meaningfully reducing the pharmacological burden (15,16,17,18), while adaptable vibrotactile stimulation achieved a 48.78% reduction of tremor. To hit a 50% total tremor reduction target, propranolol dose dropped from 320 mg to 90 mg when combined with VTS, representing a 71.9% reduction in drug dose. That lower dose also reduced the blood pressure drop from 38.1 to 18.9 mmHg, reducing hypotensive side effect risk by 19.2 mmHg (10,11). Hill curve modeling confirmed a nonlinear dose-response relationship, and VTS was most impactful at higher suppression targets.

All five vibration levels significantly reduced tremor compared to baseline, each with p < 0.001, confirmed by Kruskal-Wallis testing across conditions (p = 0.040) and bootstrap confidence interval resampling over 10,000 iterations. The optimal vibration level was 0.78 Grms, achieving a 37.3% mean reduction with a median amplitude of 2.05 versus a baseline of 3.75 g^2^, and a peak single-condition suppression of 69.0% at a vibration setting of 72 over 15 seconds. Short 15-second bursts consistently outperformed longer durations. In contrast, 90-second continuous stimulation at a vibration setting of 120 produced an 11% tremor increase, suggesting neural habituation as a limiting factor (16,18). A Friedman repeated-measures test confirmed the suppression pattern held consistently across all conditions (p = 0.0012, W = 0.320) (Figure 5).

**Figure 5.**
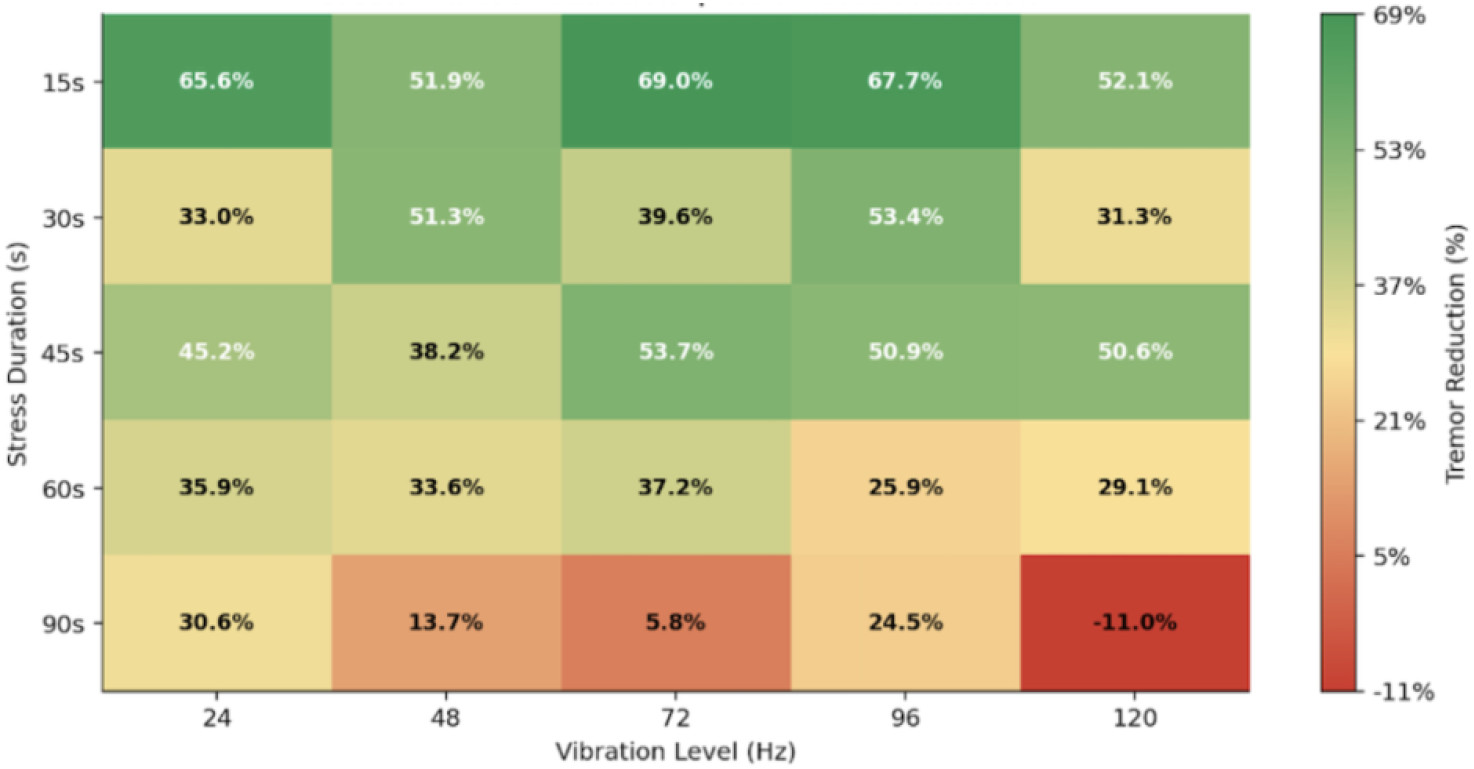
Tremor suppression increases as the vibration level increases, while also decreasing as the stress duration increases. Heatmap showing the tremor reduction across different vibration levels and stress durations (n=2). A weight was applied on the arm to recreate tremors, while vibrotactile stimulation was applied on the fingertips. Friedman test (p = 0.0012, W = 0.320).

## DISCUSSION

This project explored a novel approach to mitigate tremor in a non-monotonic way that combined stress modeling, propranolol dosage simulation, and adaptive vibrotactile stimulation instead of only relying on medication, with the goal of maximizing the effectiveness of VTS while ensuring that we minimized pharmacological side effects.

The stress-response model achieved an AUC of 0.919, showing reliability in using stress levels as an input for estimating tremor severity and dosage response. This provided the basis for quantifying propranolol’s stress-reduction effects under different conditions (3,20). Results from the simulation showed that propranolol reduced tremor amplitude, but its effectiveness followed a curve which has diminishing results. Higher doses had smaller gains while also increasing the risk of hypotension, fatigue, dizziness, and cognitive fog through low blood pressure (5,8,9,10,11).

Vibrotactile stimulation stood out as a complement to pharmacological treatment with fewer side effects. Rather than suppressing neurotransmitters, it worked by disrupting the synchronized muscle oscillations (16,17). A Wilcoxon signed-rank test confirmed that tremor reduction during stimulation was statistically significant (p < 0.001), and a Kruskal-Wallis test showed that differences across vibration levels were also significant (p = 0.04). Unlike previous devices that mainly relied on fixed vibration frequencies, our system operated in a closed loop, continuously updating tremor characteristics and adjusting vibration intensity based on the optimal regression line: y = 120x^0.485/(2.407^0.485 + x^0.485). This finding supported the hypothesis that tremor has a non-monotonic relationship with vibration, where different amplitudes varied in how effectively they disrupted oscillatory synchronization (17,18). Further analysis showed that pairing adaptive VTS with propranolol could drastically reduce the required drug dose, lowering the chance of side effects (10,11) (Figure 6).

**Figure 6.**
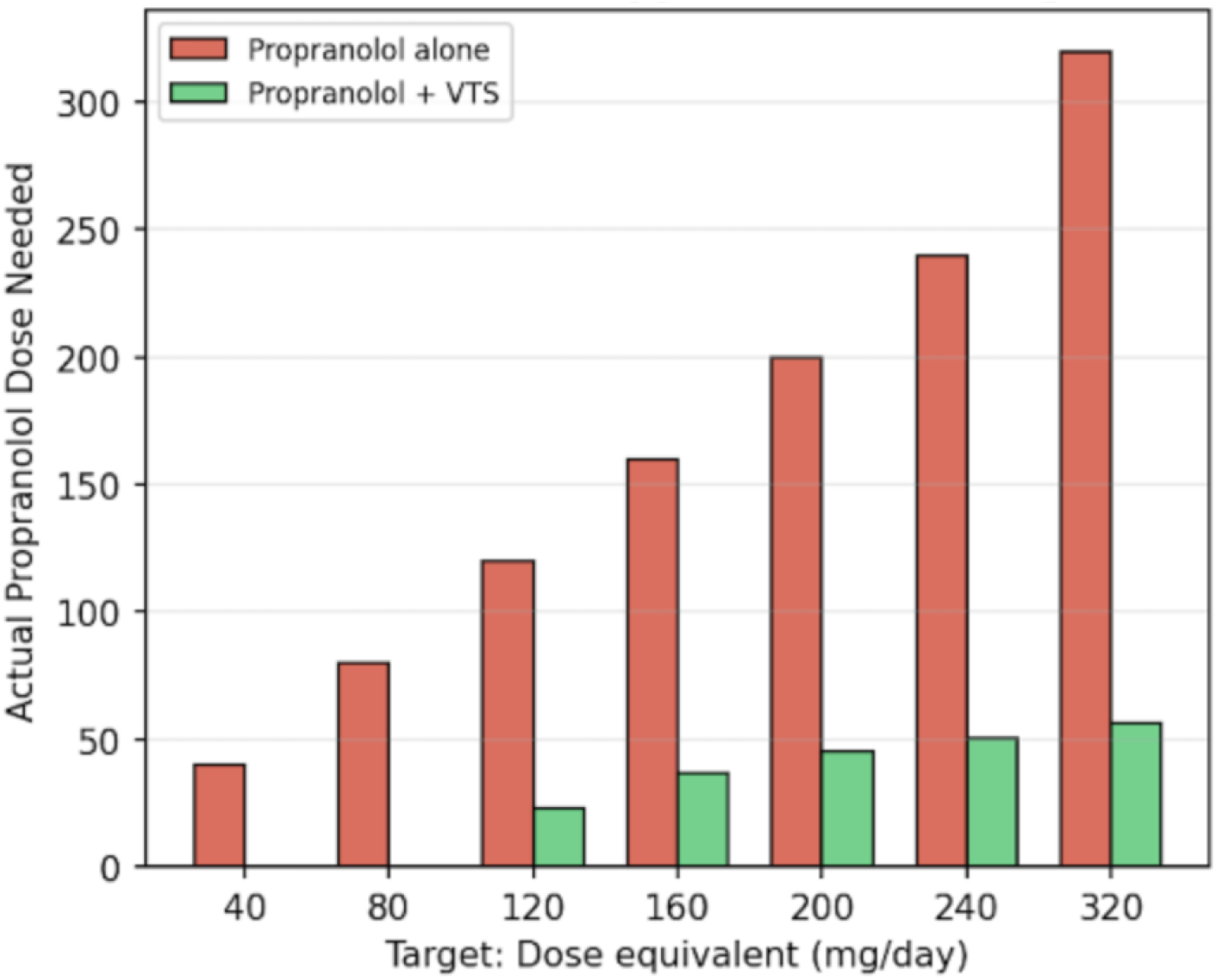
Propranolol dosage needed is reduced when using vibrotactile stimulation. Bar graph comparing modeled propranolol dose required to reach a range of target tremor-reduction levels, with and without VTS (n=2). Dosing Hill curve generated from simulated dose-response data and compared with tremor reduction achieved through VTS.

The study had several limitations. First, we used a small sample size of two subjects with 25 trials each, which limited the generalizability of our findings. We computationally simulated propranolol effects rather than being able to validate them, and we tested healthy subjects with replicated physiological tremor rather than essential tremor patients. Our trials were also short, capped at 90 seconds, so long-term habituation effects remain unknown. A wider range of vibration frequencies would also help produce a more accurate regression curve. In future work, we plan to conduct more trials in senior centers with essential tremor patients. Participants will sign consent forms, and we will also test trials with varying stimulation durations.

## MATERIALS AND METHODS

The materials utilized for this device were as follows:

- I2C DRV2605L Haptic Motor Drive Controller Module For Buzzer Vibration Motor | Quantity: 2
- MPU-6050 MPU6050 6DOF 3 Axis Gyroscope+Accelerometer Module | Quantity: 1
- ATmega328P Nano Type-C Controller Board Soldered | Quantity: 2
- Wrist Brace Support Sports Band Wrap Adjustable Carpal Tunnel Bandage | Quantity: 1
- VIBRATION LRA MOTOR 235HZ 1G | Quantity: 2
- VIBRATION ERM MOTOR 3VDC | Quantity: 4
- JUMPER WIRE M TO M 6” 28AWG | Quantity: 2
- BREADBOARD-1 | Quantity: 1

Two coin linear resonant actuators (LRAs) were used and were placed on the fingertips to deliver vibrotactile stimulation (13,15,16,22). Motion data related to tremors, including amplitude and dominant frequency, were captured by an inertial measurement unit (IMU), while real-time processing and updating were performed by an Arduino microcontroller, with vibration frequency and amplitude being controlled through a motor driver based on continuous sensor feedback.

The system was operated in a closed loop: tremor amplitude was continuously monitored by the IMU, and stimulation parameters were adjusted based on that data. The goal was to identify vibration settings that reduced tremor amplitude without also directly interfering with daily tasks (Figure 7).

**Figure 7.**
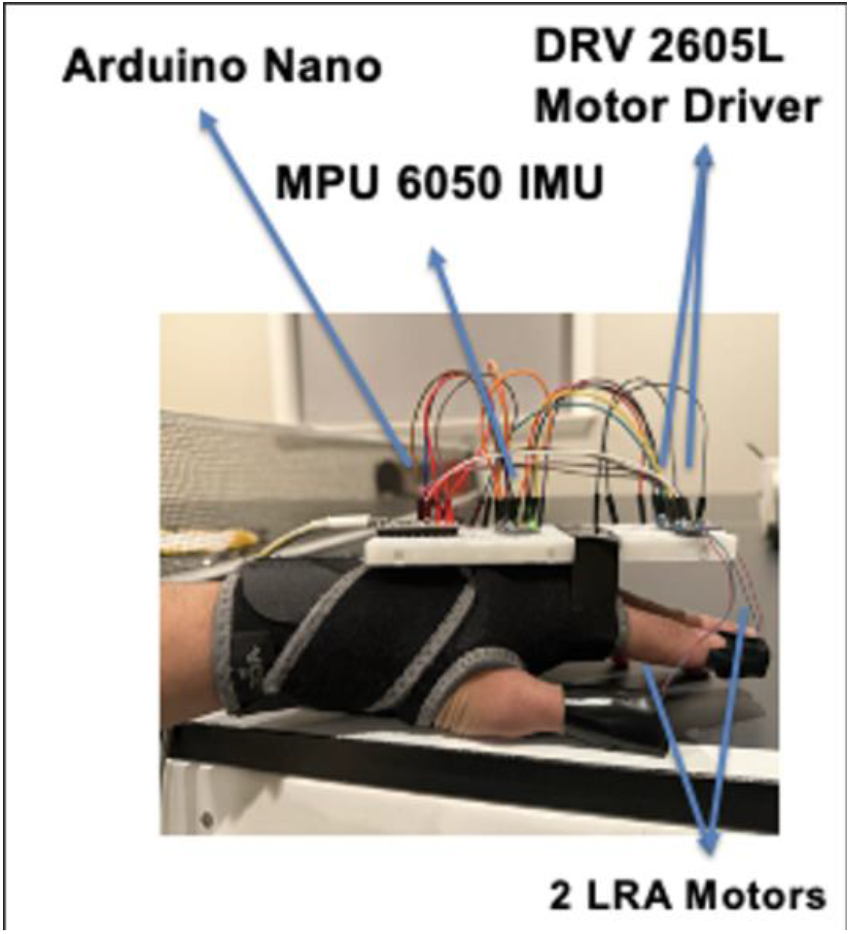
Prototype of the closed-loop vibrotactile stimulation device. Photograph of the wrist-worn prototype showing the Arduino Nano microcontroller, DRV2605L haptic motor driver, MPU-6050 inertial measurement unit, and the two linear resonant actuator (LRA) motors positioned on the fingertips.

For signal processing, the raw gyroscope output was decomposed using a db4 wavelet at five levels (Figure 8). The s5 component (0–1.6 Hz) was removed to eliminate slow postural drift and gravitational wavelets from the wrist sensor, and the d1 component was removed to filter out LRA motor buzz during active stimulation. The d2–d5 components were kept, reconstructing a signal that used only the wavelets within the 4–12 Hz tremor band range (2). A wavelet approach was chosen over FFT because tremor isn’t stationary and changes, and its frequency content shifted throughout different phases, requiring decomposition based on time frequency rather than static frequency decomposition (2,23).

**Figure 8.**
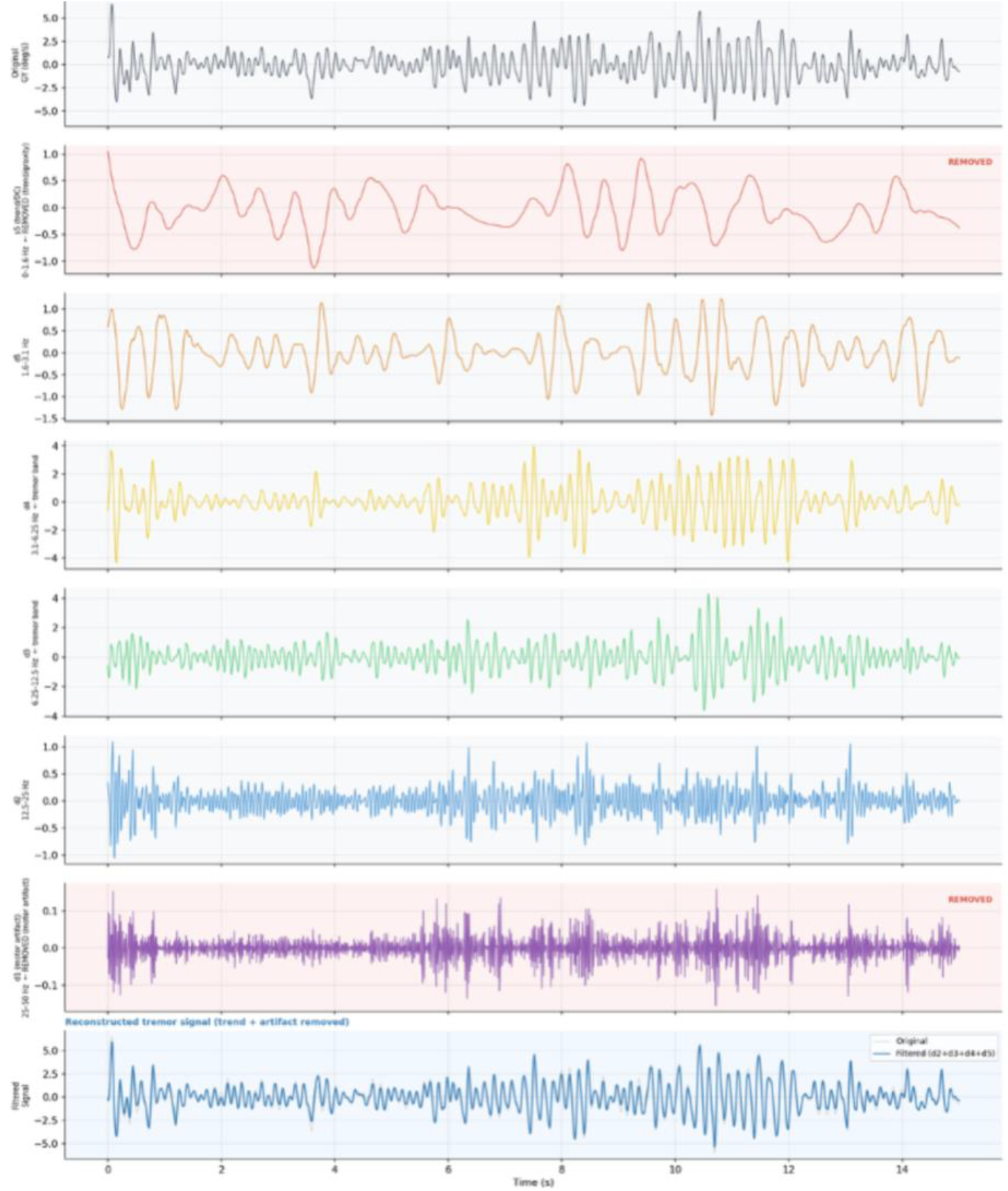
Wavelet decomposition and reconstruction of the tremor signal using a db4 wavelet function. The graph shows the raw gyroscope signal decomposed into its component frequency bands, with postural-drift and motor-artifact components removed, and the reconstructed tremor-band signal shown at bottom. Generated using Daubechies wavelet decomposition of IMU data captured from the device.

To test the device, a weight was attached to the upper arm while the device was worn on the wrist, with LRA motors placed on the thumb and index fingertips to target Pacinian corpuscle sensitivity (24). A 90-second baseline tremor was recorded with no vibration before any trials began. Trials were then run at durations of 15, 30, 45, 60, and 90 seconds using PWM values of 24, 48, 72, 96, and 120 in randomized order. A total of 50 trials were completed, with 25 per subject, and 20-minute rest periods were used between trials to reduce fatigue effects. Wavelet decomposition was applied to each trial in order to get tremor amplitude, frequency, and approximate entropy, with all data saved to Excel for more analysis.

## ACKNOWLEDGMENTS

We thank Dr. Jacob George from the University of Utah NeuroRobotics department for his help on setting our project direction.

## DATA AVAILABILITY

The data and analysis code that support the findings of this study are available from the corresponding author upon reasonable request.

## REFERENCES

1. “Differences in the Experience of Active and Sham Transcranial Direct Current Stimulation.” Kessler, S. K., et al. Brain Stimulation, vol. 5, no. 2, 2012, pp. 155–162. 10.1016/j.brs.2011.02.007.

2. Schmidt, P., et al. “Introducing WESAD, a Multimodal Dataset for Wearable Stress and Affect Detection.” Proceedings of the 20th ACM International Conference on Multimodal Interaction, 2018, pp. 400–408. 10.1145/3242969.3242985.

3. Ometov, A., et al. “Stress and Emotion Open Access Data: A Review on Datasets, Modalities, Methods, Challenges, and Future Research Perspectives.” Journal of Healthcare Informatics Research, vol. 9, no. 3, 2025, pp. 247–279. https://pmc.ncbi.nlm.nih.gov/articles/PMC12290141/.

4. Bremner, J. D., et al. “Application of Noninvasive Vagal Nerve Stimulation to Stress-Related Psychiatric Disorders.” Journal of Personalized Medicine, vol. 10, no. 3, 2020. 10.3390/jpm10030119.

5. “Practical Guide for Feature Engineering of Time Series Data.” dotData, https://dotdata.com/blog/practical-guide-for-feature-engineering-of-time-series-data/. Accessed 18 Mar. 2026.

6. Dai, D., et al. “Comparative Effectiveness of Transcutaneous Afferent Patterned Stimulation Therapy for Essential Tremor: A Randomized Pragmatic Clinical Trial.” Tremor and Other Hyperkinetic Movements, vol. 13, 2023. https://pmc.ncbi.nlm.nih.gov/articles/PMC10588491/.

7. Chahine, L., et al. “Medical Devices for Tremor Suppression: Current Status and Future Directions.” Biosensors, vol. 11, no. 4, 2021. https://pmc.ncbi.nlm.nih.gov/articles/PMC8065649/.

8. Bhidayasiri, R., et al. “The Evolution of Therapeutic Strategies in Essential Tremor: Past, Present, and Future.” Tremor and Other Hyperkinetic Movements, vol. 11, 2021. https://pmc.ncbi.nlm.nih.gov/articles/PMC8015902/.

9. van der Heide, Anouk, et al. “Propranolol Reduces Parkinson’s Tremor and Inhibits Tremor-Related Activity in the Motor Cortex: A Placebo-Controlled Crossover Trial.” Annals of Neurology, vol. 97, no. 4, Dec. 2024. 10.1002/ana.27159.

10. Bhidayasiri, R., and D. Tarsy. “Does Propranolol Reduce Parkinson-Related Tremor?” NEJM Journal Watch, 25 Feb. 2025, https://www.jwatch.org/na58448/2025/02/25/does-propranolol-reduce-parkinson-related-tremor.

11. Saporta, Raphaël, et al. “Simulation-Based Evaluation of the Impact of Dose Fractionation Study Design on Antibiotic PKPD Analyses.” JAC-Antimicrobial Resistance, vol. 7, no. 2, Mar. 2025. 10.1093/jacamr/dlaf057. Accessed 27 May 2026.

12. Herndon, D. N., et al. “Propranolol Decreases Cardiac Work in a Dose-Dependent Manner in Severely Burned Children.” Annals of Surgery, 2010. https://pmc.ncbi.nlm.nih.gov/articles/PMC3008513/.

13. Koller, W. C., and V. L. Royse. “Propranolol and Propranolol-LA in Essential Tremor: A Double Blind Comparative Study.” Journal of Neurology, Neurosurgery & Psychiatry, vol. 49, no. 6, 1986, pp. 692–695. 10.1136/jnnp.49.6.692.

14. Koller, W. C. “Dose-Response Relationship of Propranolol in the Treatment of Essential Tremor.” Archives of Neurology, vol. 43, no. 1, Jan. 1986, pp. 42–43. 10.1001/archneur.1986.00520010038018.

15. Elias, W. J., et al. “Local Vibrational Therapy for Essential Tremor Reduction: A Clinical Study.” Tremor and Other Hyperkinetic Movements, vol. 10, 2020. https://pmc.ncbi.nlm.nih.gov/articles/PMC7589646/.

16. Pfeifer, K. J., et al. “Coordinated Reset Vibrotactile Stimulation Induces Sustained Cumulative Benefits in Parkinson’s Disease.” Frontiers in Physiology, vol. 12, 2021. 10.3389/fphys.2021.624317.

17. Liu, W., T. Kai, and K. Kiguchi. “Tremor Suppression With Mechanical Vibration Stimulation.” IEEE Access, vol. 8, 2020, pp. 226199–226212. 10.1109/ACCESS.2020.3045023.

18. Oh, V. M., et al. “Effects of Long-Acting Propranolol on Blood Pressure and Heart Rate in Hypertensive Chinese.” British Journal of Clinical Pharmacology, vol. 20, no. 2, Aug. 1985, pp. 144–147. 10.1111/j.1365-2125.1985.tb05046.x.

19. Shand, D. G. “Propranolol.” British Journal of Clinical Pharmacology, vol. 4, no. 1, 1977, pp. 3–7. 10.1111/j.1365-2125.1977.tb00660.x.

20. Lora-Millan, Julio S., et al. “A Review on Wearable Technologies for Tremor Suppression.” Frontiers in Neurology, vol. 12, Aug. 2021. 10.3389/fneur.2021.700600.

21. “Sensing Vibration.” Harvard Brain Science Initiative, Harvard University, https://brain.harvard.edu/hbi_news/sensing-vibration/. Accessed 18 Mar. 2026.

22. Cabral, Ariana Moura, et al. “On the Effect of Vibrotactile Stimulation in Essential Tremor.” Healthcare, vol. 12, no. 4, Jan. 2024, p. 448. 10.3390/healthcare12040448.

23. Shah, Manthan. “Non-Invasive Passive Hand Tremor Control Orthoses.” University of British Columbia, Dec. 2022, https://open.library.ubc.ca/media/stream/pdf/24/1.0422477/4.

24. Riviere, C. N., et al. “Simulation of Tremor.” Neurological Research, 2002. https://www.sciencedirect.com/science/article/pii/S0987705302002964.

